# Revealing the functions of supra-temporal and insular auditory responsive areas in humans

**DOI:** 10.1101/2020.03.30.015289

**Authors:** Qian Wang, Lu Luo, Na Xu, Jing Wang, Yayue Gao, Siqi Li, Mengyang Wang, Pengfei Teng, Yuguang Guan, Jian Zhou, Tianfu Li, Xing Tian, Guoming Luan

## Abstract

The human auditory sensory area, which includes primary and non-primary auditory cortices, has been considered to locate in the supra-temporal lobe for more than a century. Recently, accumulating evidence shows that the posterior part of insula responses to sounds under non-task states with relevant short latencies. However, whether posterior insula (InsP) contribute to forming auditory sensation remains unclear. Here we addressed this issue by recording and stimulation directly on the supra-temporal and insular areas via intracranial electrodes from 53 epileptic patients. During passive listening to a non-speech sound, the high-γ (60-140 Hz) active rate of InsP (68.8%) was approximate to the non-primary auditory areas (72.4% and 79.0%). Moreover, we could not distinguish InsP from supra-temporal subareas by either activation, latency, temporal pattern or lateral dominance of sound induce high-γ. On the contrary, direct electrical stimulation evoked auditory sensations effectively on supra-temporal subareas (> 65%), while sparsely on InsP (9.49%). The results of cortico-cortical evoked potentials (CCEPs) showed strong bidirectional connectivity within supra-temporal areas, but weak connectivity between supra-temporal areas and InsP. These findings suggest that even the InsP has similar basic auditory response properties to the primary or non-primary cortex, it may not directly participate in the formation of auditory perception.

## INTRODUCTION

Unlike the visual cortex, the hierarchical organization of the human auditory cortex is not fully investigated. Flechsig (1908) first identified supra-temporal gyrus as the auditory cortex in humans, who also observed that Heschl’s gyrus (HG) receives denser thalamic inputs from the medial geniculate body. Nowadays, HG is regularly selected as an anatomic marker for the primary auditory cortex while the rest of the supra-temporal lobe is defined as the non-primary cortex (Poeppel et al., 2012).

Is the supra-temporal area in charge of all the early auditory processing? This hypothesis is challenged by the findings of auditory-responsive neurons in non-human primates’ insula (Sudakov et al., 1971; Bieser, 1998; Bieser and Müller-Preuss, 1996; Remedios et al., 2009). Using high-density electrodes arrays, Remedios and colleges (2009) had found that 69% of the insular neurons (clustered in the caudal part) respond to pure tones, with longer averaged response latencies than auditory cortex neurons. These findings support the fMRI activations of the insula during simple and complex auditory paradigm in human participants (Schirmer et al., 2012; Herdener et al.,2008; Zatorre et al., 1994; Meyer et al., 2002; Kotz et al., 2003; Wong et al., 2004; Sander and Scheich, 2005). However, the understanding of the basic auditory response properties of the human insula is larger unknown.

Lying deeply inside the Sylvain fissure, the insula is difficult to reach for stander non-invasive electrophysiological technologies in human participants. In that case, intracranial electroencephalogram (iEEG) monitoring in epileptic patients provides a unique technology that could record the insular signals directly with both high spatial and temporal resolutions. Recently, accumulating intracranial evidence shows that insula could respond to acoustic stimuli with relatively short latencies (Cui et al., 2017; Zhang et al., 2018; Woolnough et al., 2019). Especially, Woolnough and colleges (2019) found that the activation patterns of posterior insular were like those of superior temporal gyrus (STG). On the other hand, direct electrical stimulations on some of the sites located in the human insula (clustered posteriorly) could induce auditory sensory (Mazzola et al., 2017). This evidence motivated us to ask a question: whether the posterior insula act as a part of the auditory sensory cortex?

To answer that, here we performed direct recordings and stimulations via the invasive electrodes’ implanted in 53 drug-resistant epileptic patients. This large data set provided approximately full coverage of both supra-temporal and insular cortices. The supra-temporal area was further divided into HG, and the anterior, middle, and posterior parts of STG, while the insular area divided into anterior (InsA) and posterior (InsP) regions by the central insular sulcus. We systematically investigated: (1) the acoustic response properties (latency, activation rate, temporal patterns, and lateral dominance) of supra-temporal and insular cortices by recording the sound-induced high-γ activities; (2) the auditory sensation sites on supra-temporal and insular cortices by electrical brain stimulation (EBS); (3) the functional connectivity between supra-temporal and insular cortices by both cortical-cortical evoked potential (CCEP) technology and Granger causality analyses. Last but not the least, we also recorded the extracellular field potentials using micro contacts to verify the true regional activity source of the InsP.

## RESULTS

### Activation rate and latency in auditory responsive areas

In all 55 participants, 953 out of 6534 contacts which located on the Heshl’s gyrus (HG, *n* = 168), anterior superior temporal gyrus (aSTG, *n* = 82), middle superior temporal gyrus (mSTG, *n* = 105), posterior superior temporal gyrus (pSTG, *n* = 105), anterior insula (InsA, *n* = 194), and posterior insula (InsP, *n* = 299) were selected (Figure 1A and B).

For each contact, sound-induced high-γ activity (60-140 Hz) was extracted. Firstly, we grouped auditory evoked high-γ activities in ROI level to roughly estimate their responsiveness to the non-speech sound. As the waveforms showed in Figure 1B, HG, post-onset significances were found in InsP, pSTG, and mSTG, but not in InsA or aSTG (*t*-tests against the baseline, with *Bonferroni* correction). Considering the sparse organization of the cortex, we also applied the single-trial level statistical analyses. As shown in Figure 1B and C, 524 out of 953 contacts were significantly sound-responsive under bilateral sound conditions (one-sample *t*-tests against the baseline). Nearly all the contacts located in HG significantly responded to the sound (active rate = 94.6%) while the activation rates in InsP (68.8%), pSTG (72.4%), and mSTG (79.0%) were approximately similar, which were higher than those in InsA (37.6%) and aSTG (28.0%) (Figure 1C). The ranges of activation rates were similar under monaural input conditions, where the ipsilateral input condition showed generally lower activity (Figure S1). Considering both group and individual level results, only sound-responsive contacts located on HG, InsP, pSTG, and mSTG were entered further analyses.

In the group level, HG showed stronger activation (all *p* < 0.001, with *Bonferroni* adjustment) and shorter first significant latency (all *p* < 0.01, with *Bonferroni* adjustment) than InsP, pSTG, or mSTG (Figure 1D). To test whether the sound-induced high-γ conform to a certain spatial distribution in either insular or supra-temporal areas, we calculated the Pearson correlation between the contact-level activations and latencies and their MNI coordinates, respectively. As shown in Figure 1E, a high-to-low gradient for activation and an early-to-late gradient for latency along the medial-lateral axis was found on the supra-temporal plane (HG, pSTG, and mSTG) (both *p* < 0.001), consistent with previous literature (Nourski et al., 2014; Patel et al., 2018). While in the InsP, a low-to-high gradient for activation a late-to-early gradient for latency along the anterior-posterior axis was found (both *p* < 0.001), consistent with the findings in non-human primates (Remedios et al., 2009). Similar distributions could also be observed under monaural input conditions (Figure S2).

**Figure 1.**
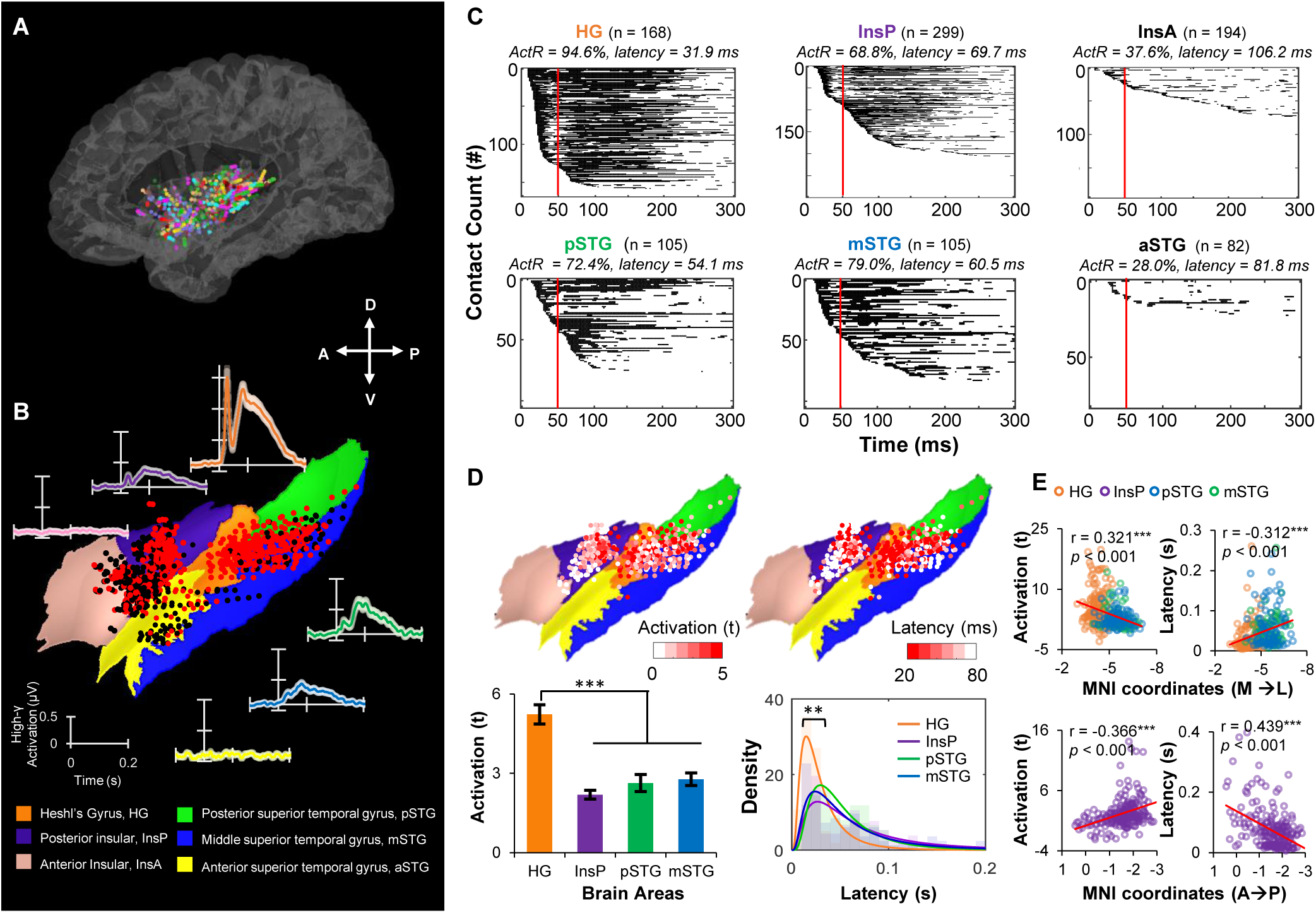
The auditory responsive regions within the human supra-temporal lobe and insula. (A) Spatial locations of selected sEEG contacts from all patients on a BCI-DNI template brain (*55 patients, 953 contacts, projected to left hemisphere*). (B) Curve diagrams show the average auditory evoked high-γ(60-140 Hz) responses within 6 regions of interest (ROIs), respectively. Shaded areas indicate standard errors. All 953 contacts are projected on the colored ROI map, where red dots represent the significant contacts and black dots represent the non-significant contacts. (C) Raster plots of the significance against the baseline in 6 ROIs. Each line represents one contact and black dots represent the significant time points against the baseline (*t*-tests in single-trial level, *Bonferroni* corrected *p* < 0.05). (D) Contacts located on HG, InsP, pSTG, and mSTG, color-coded by activation (*left top*) and latency (*right top*). Comparison of the activation (*left bottom*) and the latency (*left bottom*) among these 4 ROIs. (E) The correlation between the response properties (activation and latency) and MNI coordinates in the supra-temporal area (HG, pSTG, and mSTG) and InsP. *ActR: activation rate; A→P, from anterior to posterior; M→L, from medial to lateral. **, p < 0.01; ***, p < 0.001.*

### Unsupervised clustering for temporal response patterns

According to a previous ECoG study, the temporal patterns of the sound-induced high-γ activity in human STG could be divided into “onset” and “sustained” types (Hamilton et al., 2018). However, whether there exists a similar classification in HG and InsP, which are buried inside the lateral fissure and can be reached by ECoG, is still unclear. As shown in Figure 1B, we visually observed that the averaged high-γ waveforms might be divided into least two stages: a 0-50 ms early component, and a 50-300 ms late component (Figure 1B).

We applied an unsupervised clustering algorithm, non-negative matrix factorization (NMF), on the high-gamma time series from 524 contacts across 53 patients. Although Hamilton and colleges (2018) showed that the sound-induced high-γ in STG could be roughly divided into two classes, our sEEG contacts in the current study also covered InsP and HG, which could hardly be reached by ECoG. In the case of InsP and HG will bring new classification features, we firstly investigated the optimal number of classifications. Explanatory powers (*r* square) under different cluster number *k* from 2 to 10. When *k* = 2, although the results explained 53.9% of the variance in the data, the “early” and “late” pattern could not be effectively distinguished (Figure S3A). When *k* = 3, the results explained 60.0% (with 6.1% added) of the variance in the data while the observed 0 to 50 ms “early” pattern could be effectively divided from the “late” ones (Figure S3B). When *k* raised to 4, the explanation of the variance only raised 3.0% (Figure S3C). Therefore, as shown in Figure 2A, we finally chose to optimally cluster response profiles into three groups.

We then measured the peak latency of each feature waveform. As shown in Figure 2A, the latencies of the first and the second components were within 100 ms while that of the third one was larger than 100 ms. According to the latency ranges of the “Onset” and the “Sustained” responses in Hamilton et al., 2018, we named the three clusters into “Onset □”, “Onset □” and “Sustained”, respectively. Contacts were then clustered into these three classes according to their maximum weights (Figure 2B, Figure S4). Clustering strengths of each ROI were quantified using silhouette indices (SIs), and all the SIs were significantly larger than the chance level (all *p* < 0.001, paired *t*-tests between the real SI and the shuffled calculations) (Figure 2C).

The proportion of contacts belonged to different classes significantly differs across the 4 ROIs (*p* = 2.7×10^−5^, *chi-square* test, right panels in Figure 2A). That is, HG and InsP contributed to a large proportion of the “Onset □” clusters, while pSTG and mSTG mainly contributed to “Onset □” and “Sustained” clusters.

To examine whether the response patterns were spatially organized, *Pearson* correlation tests between the classification weights of each cluster and MNI coordinates were conducted for supra-temporal areas and InsP, respectively. For supra-temporal areas, the weight of “Onset I” significantly decreased when the contact was located more lateral (*r*_*315*_ = 0.321, *p* < 0.001, with *Bonferroni* correction). More interestingly, we found that when the coordinates changed from anterior to posterior, the “Onset □” weights increased while the “Sustained” weights decreased (both corrected *p* < 0.05). These results were in consistent with the previous findings in STG (Hamilton et al., 2018), and further suggested that “Onset □”, but not “Onset I”, is more associated with the Hamilton’ *Onset* component. These results also indicated that the “Onset I” component might be a featured response in HG.

Spatial organizations were also found in InsP. The “Onset I” weights were significantly stronger in the dorsal part while the “Onset □” weights were significantly larger in the posterior part of the InsP (both corrected *p* < 0.05).

**Figure 2.**
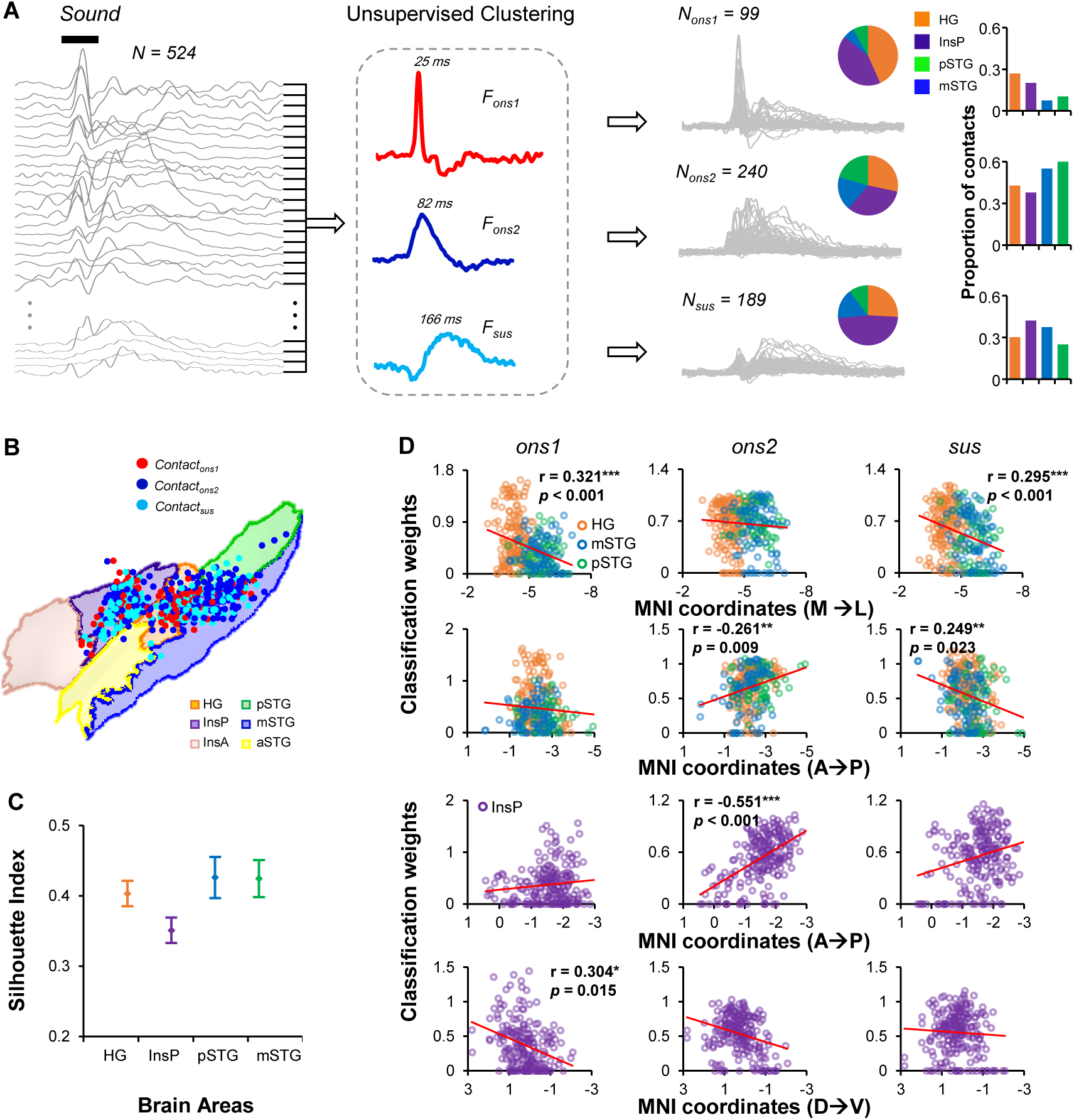
The waveforms of auditory evoked high-γactivity can be clustered into three temporally classes. (A) Illustration of the unsupervised clustering flow. Only significantly responsive contacts within HG, InsP, pSTG, and mSTG were selected to enter the clustering pool (*n* = 524). Both the featured time series of the first (*red line*, “Onset □”, *F*_*ons1*_) and that of the second cluster (*dark blue line*, “Onset □”, *F*_*ons2*_) show transient activations followed by slow inhibitions after the sound onset (both peak latencies within 100 ms), while the time series of the third cluster (*light blue line*, “sustained”, *F*_*sus*_) shows show a transient inhibition followed by slow activation instead. Contacts are then divided into three groups (*N*_*ons1*_ *= 99; N*_*ons2*_ *= 240; N*_*sus2*_ *= 189*). The proportions of the contacts in three classes are different across the 4 ROIs. (B) The distribution of the contacts from three classes. (C) The clustering goodness of fit of each brain area was tested using the silhouette index (SI). All SIs are significantly larger than the chance level (all *p* < 0.001, paired *t*-tests). Bar showed the SEM. (D) The correlation between classification weights and MNI coordinates for the supra-temporal area (HG, pSTG, and mSTG) and InsP. *A→P, from anterior to posterior; M→L, from medial to lateral; D→V, from dorsal to ventral. *, p < 0.05; **, p < 0.01; *** p < 0.001. See also Figure S3 and S4.*

### Contralateral dominance

The human auditory cortex was more responsive to sounds coming from the contralateral directions (Ackermann et al., 2011; Patel et al., 2018). However, whether the InsP also has lateral selectivity is still unclear.

Figure 3A showed a pair of symmetrical electrodes (G and GG) implanted in an individual patient (P370). In four example contacts, the sound-induced high-γ tuned to contralateral sound during both early (0-50 ms) and late (50-300 ms) periods (Figure 3B). This dominance could also be observed in the group level, that contralateral input condition had higher activations and shorter latencies in HG, pSTG, mSTG and InsP (Figure S1). To investigate the lateral dominance in contact level, paired *t*-tests were conducted between the high-γ activities under contralateral and ipsilateral input conditions, while the *t*-values were extracted as the contralateral-ipsilateral (CI) indices (Figure 3C and D). For all the auditory responsive areas, the CI indices increased when the early period turned into the late period; the proportions of contacts with significantly contralateral dominance were higher during late period (HG: 74.45%; InsP: 30.42%; pSTG: 35.71%; mSTG: 36.14%) than during the early period (HG: 32.85%; InsP: 7.92%; pSTG: 15.31%; mSTG: 12.05%) (Figure 3E and F). There also existed a small proportion of contacts significantly tuned to the ipsilateral sound during early period (HG: 4.38%; InsP: 1.67%; pSTG: 1.02%; mSTG: 3.61%) and late period (HG: 2.19%; InsP: 0.83%; pSTG: 0%; mSTG: 2.41%). For both periods, the CI indices in HG was significantly larger than those in InsP, pSTG, and mSTG (all *p* < 0.05, post hoc *t*-tests, with Bonferroni adjustments), while no significance was found among InsP, pSTG, and mSTG (all *p* > 0.1). These findings revealed contralateral preferring patterns in both the supra-temporal area and InsP, which were more effective in the late response stage.

To investigate the spatial organization of contralateral dominance, Pearson correlation tests between the CI indices and MNI coordinates were conducted for supra-temporal lobe and insula, respectively. For both time windows, the contralateral dominance decreased when the temporal contact was located more lateral (both *p* < 0.01, with *Bonferroni* correction), while the dominance increased when the insula contact was located more posterior (both *p* < 0.05).

**Figure 3.**
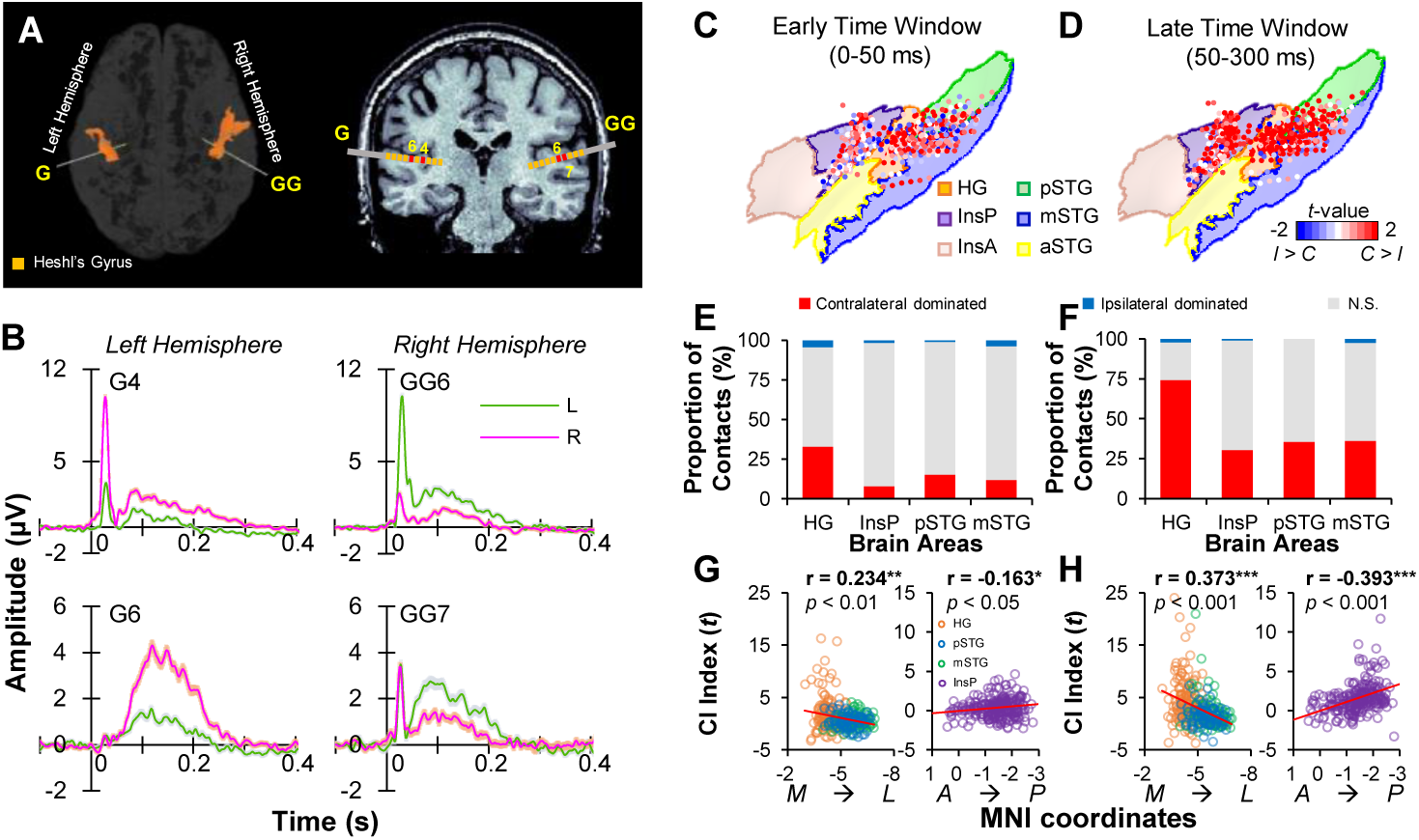
Contralateral input dominance in early and late time windows. (A) In an individual patient (P370), locations of two 10-contact sEEG electrodes implanted on the left (electrode G) and right (electrode GG) HG. (B) Averaged high-γresponses evoked by sound delivered from the left and right ear in 4 HG contacts, respectively. Contacts located on left HG (G4, G6) show preferences to right ear sound, while those located on right HG (GG6, GG7) show preferences to left ear sound, which suggesting contralateral input dominance patterns within either early (0-50 ms) or late (50-300 ms) time windows. Shaded areas indicate standard errors. (C), (E), and (G) show the distribution of the contralateral input dominance, the proportions of the significant contacts, and the correlation between CI indices and MNI coordinates within the early time window, respectively. (D), (F), and (H) show the same properties within the late time window. *CI: contralateral to ipsilateral. L, left; R, right. *, p < 0.05; **, p < 0.01; *** p < 0.001.*

### Functional mapping via electrical brain stimulation

In 42 out of 55 patients, electrical brain stimulations (EBS) were conducted to identify functional areas that needed to be considered for epileptic lobotomy plans as routine clinical tests. The contact pairs which exclusively located in HG (n = 133), InsP (n = 221), pSTG (n = 63) and mSTG (n = 71) were selected. The percentages of electrodes which related to positive sensory were different across brain areas (HG, 76.7%; InsP, 71.5%; pSTG, 42.9%; mSTG, 39.4%). The types of sensation were further classified into 8 categories (*auditory suppression, auditory hallucination, noxious sensation, somatosensory sensation, epigastria sensation, vestibular response, speech arrest, illusion*) by an experienced neuropsychologist.

Among all the contact pairs with positive EBS induced-sensory, auditory sensation and somatosensory sensation were the most common symptoms (Figure 4A and B). As shown in Figure 4A, EBS delivered to the supra-temporal areas mostly evoked auditory hallucinations (HG: 78.4%, pSTG: 66.7%, mSTG: 75.0%), while only a little proportion of EBS delivered to the InsP evoked auditory hallucinations (9.49%) (*chi-square* = 149.84, *p* = 2.85×10^−32^). By contrary, EBS delivered to the InsP areas mostly evoked somatosensory sensations (69.0%), while only a little proportion of EBS delivered to the temporal auditory areas evoked somatosensory sensations (HG: 5.9%, pSTG: 11.1%, mSTG: 17.9%) (*chi-square* = 117.72, *p* = 2.4×10^−25^). These results suggested that, although the InsP could not be distinguished from the supra-temporal areas by its auditory response properties, it may play a different role in auditory perception formation.

To further analysis the key factors of auditory perception formation, the auditory response properties of the contacts corresponded to or not to auditory sensation were divided. As shown in Figure 4C and D, the contact pairs with auditory sensation showed shorter latency (supra-temporal: *t*_*231*_ = -2.576, *p* = 0.011; InsP: *t*_*206*_ = -2.169, *p* = 0.026, independent *t*-tests) and higher activation (supra-temporal: *t*_*231*_ = 1.583, *p* = 0.115; InsP: *t*_*206*_ = 4.054, *p* = 7.1×10^−6^, independent *t*-tests) than those with other sensations.

In both supra-temporal areas and InsP, most of the auditory sensation was reported from the contralateral side (73.0%). Therefore, we further classification the contact pairs with auditory sensation into two groups, one with contralateral sound sensation, and the other-with the sensation in other locations. Independent *t*-tests showed that the CI indices of contralateral sensation contacts were significantly higher than those of other location contacts during the sustained period (50 to 300 ms) (*t*_*100*_ = 3.23, *p* = 0.002). No significance was found during the onset period (*t*_*100*_ = 1.50, *p* = 0.137).

In sum, we found that EBS delivered on the auditory responsive sites in InsP could only induce a small proportion of auditory sensation. On the other hand, our results further suggested that the sites which contributed to auditory sensation may be related to their high-γ responsive patterns but not strictly related to the brain areas they belong to.

**Figure 4.**
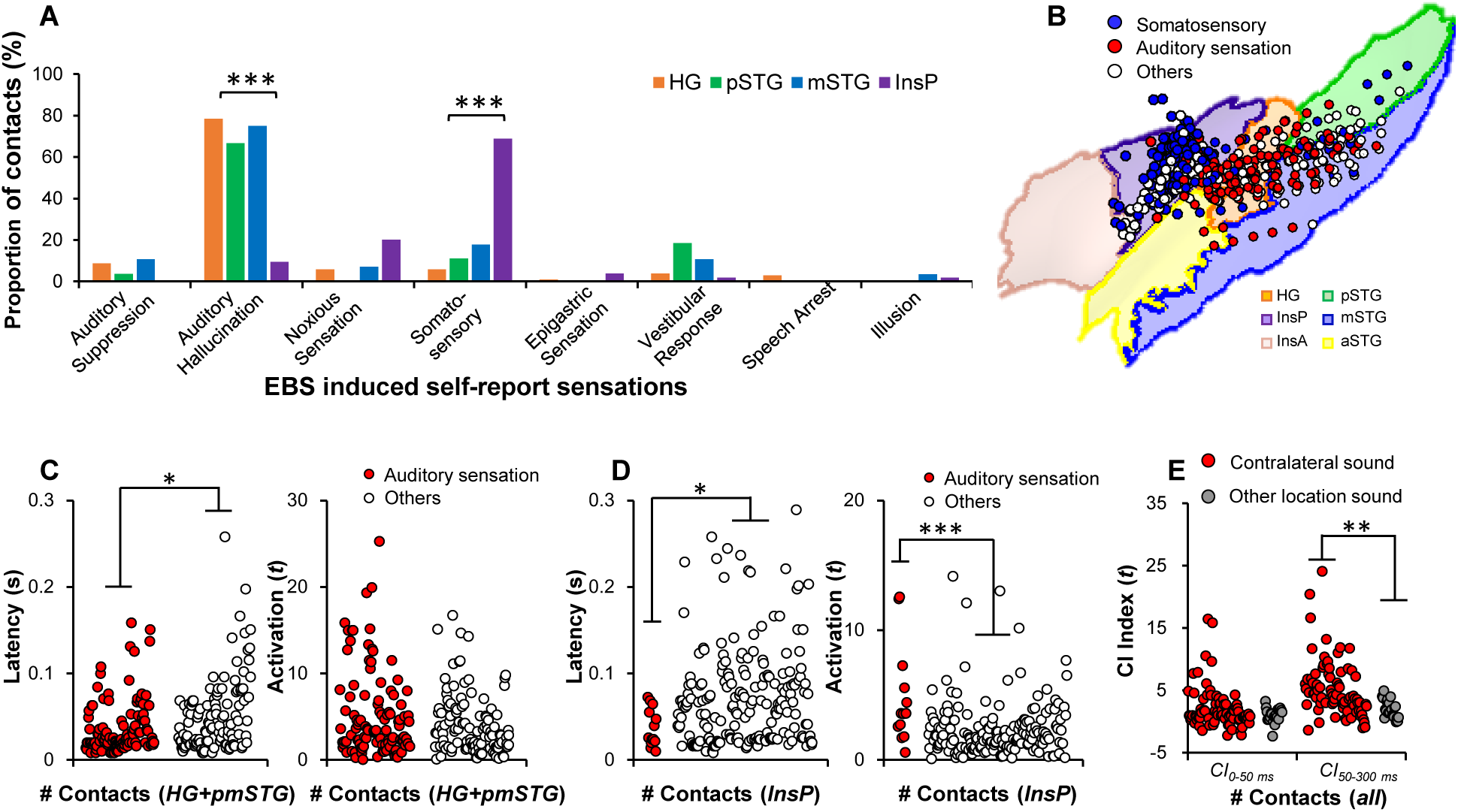
Functional mapping results revealed by the electrical brain stimulation (EBS). (A) The proportion of contacts with different EBS induced sensations across 4 ROIs. EBS delivered on HG, mSTG and pSTG contacts mainly induced auditory sensation (hallucination and suppression), while those delivered on InsP contacts mainly caused the somatosensory sensation. (B) Distributions of contacts with auditory or somatosensory sensations. (C) Comparisons of the latency and the activation of the contacts with or without auditory sensation in the supra-temporal area (HG, pSTG, and mSTG). (D) Comparisons in InsP. (E) Comparisons of the CI index (*CI*_*0-50 ms*_ *and CI*_*50-300 ms*_) of the contacts with auditory sensation from contralateral or other locations. *\*, p < 0.05; **, p < 0.01; ***, p < 0.001.*

### Connectivity between InsP and supra-temporal areas

The results of EBS implied that the auditory responsive sites in InsP may be dissociated from the auditory sensation system. Therefore, we conducted both cortico-cortical evoked potential (CCEP) and Granger causality analysis to investigate the functional connectivity among the InsP and supra-temporal areas.

CCEP was conducted in 5 patients (P359, P409, P420, P421, and P422). In the individual patient level, single-pulse electrical stimulation delivered on HG contacts (Figure 5A) could significantly evoke potentials in HG (Figure 5B) but not in InsP (Figure 5C), while electrical stimulation delivered in InsP contacts (Figure 5D) could significantly evoke potentials in InsP (Figure 5F) but not in HG (Figure 5E). These observations suggested weak connections between InsP and HG. In the group level, the CCEP amplitudes between HG and pSTG/mSTG were significantly higher than those between HG and InsP (both *p* < 0.001, post-hoc *t*-tests) (Figure 5H). Furthermore, the results of Granger causalities of the sound-induced high-γ activities also supported the stronger connection between HG and pSTG/mSTG than those between HG and InsP (both *p* < 0.001, post-hoc *t*-tests) (Figure 5I).

**Figure 5.**
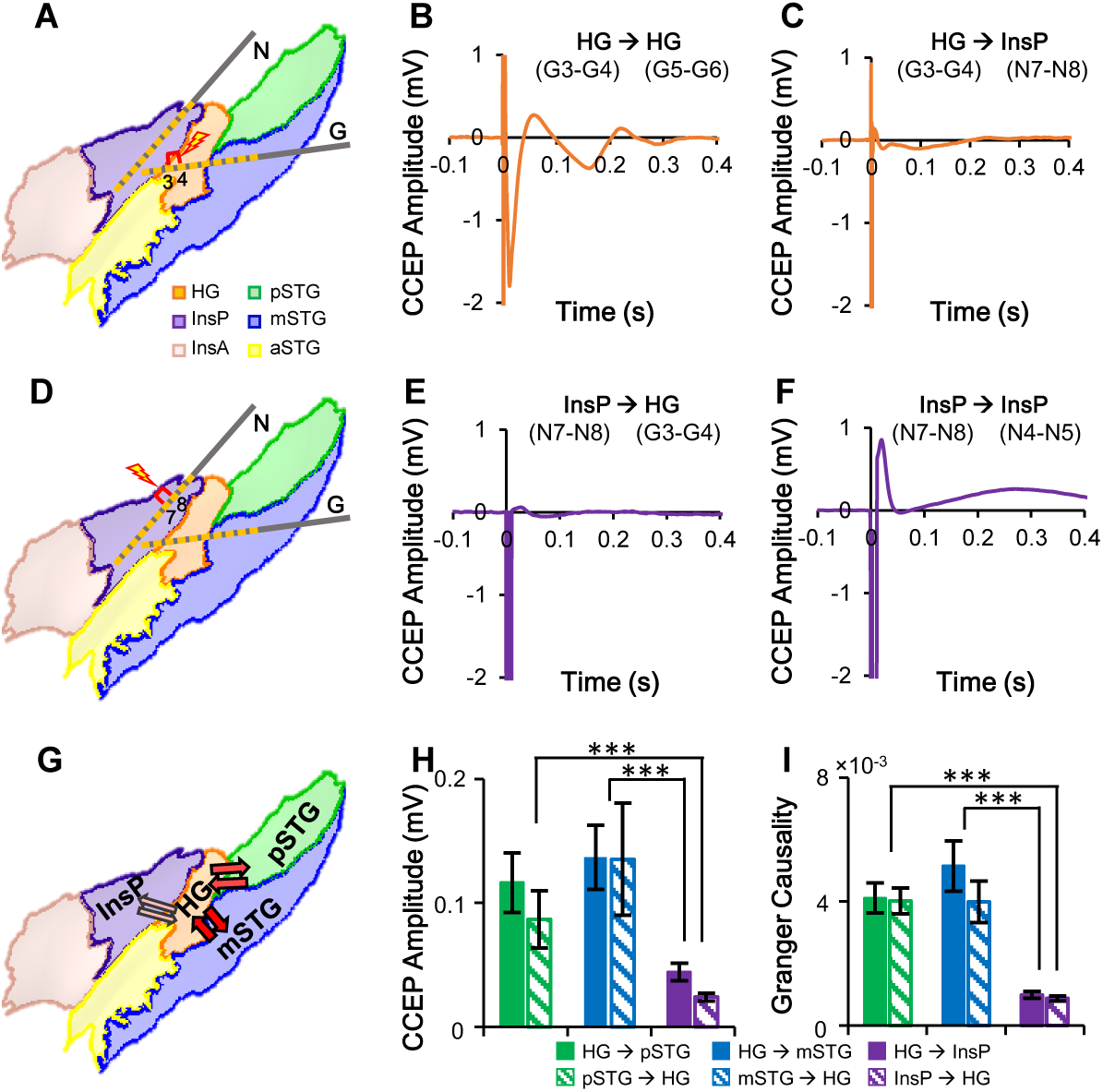
HG shows strong connections with pSTG/mSTG but a weak connection with InsP. In an individual patient (P422), the single pulse of electrical stimulation on HG (G3-G4, *panel A*) could evoke a remarkable cortical-cortical evoked potential (CCEP) on HG (G5-G6, *panel B*), but not on InsP (N7-N8, *panel C*). On the other hand, the electrical stimulation on InsP (N7-N8, *panel D*) could evoke a significant CCEP on InP (N4-N5, *panel F*), but not on HG (G3-G4, *panel E*). (G), (H) and (I) show that both the group averaged CCEPs (4 patients) and the Granger causalities of evoked high-γ (55 patients) revealed strong connections between HG and pSTG/mSTG, but a weak connection between HG and InsP. *\*\*\*, p < 0.001.*

**Figure 6.**
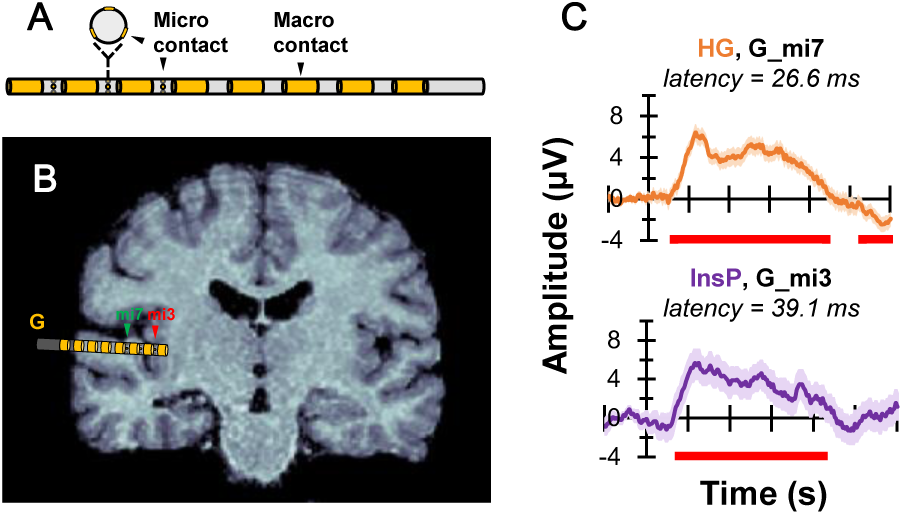
Recording auditory evoked extracellular potential in HG and InsP by micro-contacts in patient P374. (A) Illustration of micro and macro contacts on a hybrid depth electrode. (B) Location of the hybrid macro-micro electrode G implanted on the patient’s individual MR scans. For the micro contacts, the *mi7* was located on HG while *mi3* was located on InsP. (C) Auditory evoked potential in HG (mi7) and InsP (mi3) are both significant. Shaded areas indicate standard errors. Red bars below the *x*-axis represent the periods of significance (single-trial level *t*-test, *p* < 0.05, with *Bonferroni* corrections).

### Local signals in insula: evidence from micro contacts

The question may arise that whether the auditory evoked potential in InsP was a local signal or was just spread from the HG. In a rare opportunity when hybrid macro-micro electrodes were implanted in a 46 years old female patient, this speculation was tested by analyzed the auditory evoked potential in micro contacts. Totally 4 macro-micro electrodes were implanted and one was located at the junction between InsP and HG (Figure 6). As shown in Figure 6C, both micro contacts located at InsP (mi3) and HG (mi7) were significantly active than the baseline using single-trial *t*-tests (*p* < 0.05 with *Bonferroni* adjustments). The first significant latency of InsP was 39.1 ms, and that of HG was 26.6 ms, both within the ranges from the macro contact in the current study. These results confirmed that the early auditory response in InsP was a local signal, rather than the propagation from neighboring HG.

## DISCUSSION

In the current study, we applied multiple intracranial approaches including event-related high-γ activity, direct brain stimulation, and functional connectivity analysis to investigate whether the InsP could be defined as the part of the auditory sensory cortex. By analyzing the first significant latency, activation rate, temporal patterns, and lateral dominance of the sound-induced high-γ activities, we could not distinguish the InsP from the subareas of the supra-temporal lobe. However, EBS applied to the auditory responsive InsP sites mostly evoked somatosensory sensation and rarely evoked auditory sensation. On the other hand, CCEP and Granger causality analyses consistently showed weak connectivity between InsP and supra-temporal subareas. Together, our findings argue that InsP might be functionally isolated from the auditory sensory system.

### Basic auditory response properties in the supra-temporal area and InsP

The auditory responsive supra-temporal area in humans could be divided into multiple fields base on the anatomical markers (Hackett et al., 2001; Morosan et al., 2001; Rademacher et al., 1993; Wallace et al., 2002; Wessinger et al., 2001). There may exist a hierarchical organization in the auditory cortical system, where the major part of the HG is marked as the primary auditory cortex (River and Clarke, 1997). In the current study, we divided the supra-temporal lobe into 4 subregions: HG, aSTG, mSTG, and pSTG (Joshi et al., 2017). Under the passive listening condition, the aSTG responded sparsely to the non-speech sound. Thus, we defined HG, mSTG, and pSTG as the auditory responsive supra-temporal areas (Nourski et al., 2014; O’Sullivan et al., 2019; Patel et al., 2018).

According to previous intracranial recordings in humans, the primary auditory area is generally higher in activation and shorter in latency than the non-primary auditory area (Nourski et al., 2014; O’Sullivan et al., 2019; Patel et al., 2018). Our data confirmed this organization by showing HG higher in activation and shorter in latency than either mSTG or pSTG. No difference was found between mSTG and STG (Figure 1C). Thus, we equaled HG as the primary auditory area and grouped mSTG and pSTG as the non-primary auditory are, to provide comparing templates for the auditory responsive insular.

Our data showed that InsA only responded to sound sparsely and failed to reach the significance level in the group level (Figure 1B). On the contrary, 68.8% of contacts located in InsP robustly responded to the non-speech sound (Figure 1C) and reached the significance level in the group level. These findings are consistent with the previous intracranial studies (Zhang et al., 2018; Woolnough et al., 2019), which could confirm the observation of InsP activation during the simple and complex auditory task in non-invasive studies (Schirmer et al., 2012; Herdener et al., 2008; Zatorre et al., 1994; Rumsey et al., 1997; Meyer et al., 2002; Kotz et al., 2003; Wong et al., 2004; Sander and Scheich, 2005).

Using the large sEEG dataset in the current study, we provided full coverage of human InsP (*n*_*patient*_ = 54; *n*_*contact*_ = 299), which created a credible comparison between InsP and supra-temporal subareas. Our data showed that the activation, latency (Figure 1C and D) and lateral selectivity (Figure 3E and F) of InsP are close to the non-primary auditory areas. On the other hand, these three properties show coherent spatial organizations; the contacts with higher activations, shorter latencies, and stronger contralateral input dominance generally clustered in the posterior part of the InsP (Figure 1E; Figure 3G, H), which is comparable with the findings in non-human primates (Remedios et al., 2009).

The auditory cortex encodes various temporal information (Brasselet et al., 2012; Fishman and Steinschneider, 2009; Ince et al., 2013; Malone et al., 2010,). This capability may benefit from the separated neuron populations which selectively respond to different temporal features (Malone et al., 2015; Hamilton et al., 2018). One of the lesion studies suggested that InsP may involve in auditory temporal processing (Bamiou et al., 2006). However, single-unit recording in non-human primates found most insular neurons fail to follow the temporal structure of the tone trains. To investigate the temporal response pattern of InsP, and compare it with the primary and non-primary auditory area, we conducted an unsupervised clustering algorithm to extract the features among all the auditory responsive contacts. Three temporal features were selected: “Onset □”, “Onset □”, and “Sustained” as shown in Figure 2A, InsP didn’t show a unique temporal pattern but rather contained all three temporal patterns. To be noted, the “Onset □” type equals the “Onset” type in Hamilton et al., 2018, which characters in its posterior distribution in the supra-temporal plane (Figure 2B, D). Interestingly, the contact proportions of mSTG (7.5%) and pSTG (10.5%) classed into “Onset I” type were smaller than those of HG (27%) and InsP (20.1%). We then found that the “Onset I” type contacts mainly clustered on HG (medial to mSTG and pSTG) and the dorsal part of InsP. One of the reasonable explanations is that the “Onset I” is a featured response pattern in HG and dorsal InsP. It could also answer us why the “Onset I” could not be extracted from Hamilton’s (2018) dataset which mainly contained ECoG sites on lateral STG, leaving the “Onset II” type being comparable to the Hamilton’s “Onset” type.

Together, InsP could not be distinguished from the supra-temporal areas based only on their auditory response properties. However, to tested whether InsP could be defined as a part of non-primary auditory cortices, we need more direct evidence.

### Forming the auditory perception or not

The most ancient and direct approach to test whether a local area in the human cerebral cortex corresponded to a certain function is to effectively stimulate it *in vivo*, with the simultaneous observation of behavioral changes (Penfield and Rasmussen, 1950; Penfield and Jasper, 1954). Up to now, EBS becomes a routine clinical practice to map the functional involvements in sensation, motor, and other cognitive functions in epileptic patients with intracranial electrodes implantations (Selimbeyoglu and Parvizi, 2010). Major stimulations in the supra-temporal structures were associated with auditory hallucinations, or perceptual changes of sound quality from the real environment (Penfield and Perot, 1963; Fenoy et al., 2006), while the stimulation of InsP was mostly associated with somatosensory (Isnard, 2004; Nguyen et al., 2009; Afif et al., 2010; Stephani et al., 2010; Pugnaghi et al., 2011; Mazzola et al., 2017). Recently, Mazzola and colleges (2017) reported 550 clinical responses evoked by 669 insula stimulations, among which a small proportion of stimulations results in self-report auditory sensation.

One of the plausible explanations is that InsP contains both the somatosensory region and the auditory region. However, this hypothesis was questioned by the findings in the current study. We found that EBS delivered to the InsP evoked 69.0% somatosensory sensations and 9.5% auditory hallucinations, while the activation rate of InsP during passive listening condition was 68.8%. As shown in Figure 4, although the auditory sensation related contacts had shorter latencies and higher activations than other contacts, there is still a considerably large proportion of auditory responsive InsP contacts which correspond to the somatosensory sensation when EBS was delivered. This finding doesn’t support the idea that the InsP is an auditory area. Therefore, an alternative perspective is that InsP monitors the information from the auditory system and acts as a role in auditory and somatosensory integration (Rodgers et al., 2008; Woolnough et al., 2019).

Another way to test whether the InsP is an auditory area is to examine the connection between InsP and auditory. Tracing studies in non-human primates suggested possible connectivity between InsP and thalamus (Burton and Jones, 1976; Jones and Burton, 1976) and HG (Pandya et al., 1971; Augustine, 1985; Mesulam and Mufson, 1985; Pandya and Rosene, 1985; Cauda et al. 2011; Cerliani et al. 2012; Cloutman et al. 2012; Dennis et al. 2014). Using resting-state or task functional magnetic resonance imaging (fMRI) revealed that InsP connects to the auditory cortex (Nomi et al., 2016; Zhang et al., 2018) the auditory thalamus (Glasser et al., 2016). However, both the CCEP data and Granger causality results in the current studies showed that the effective connectivity between InsP and HG is much weaker than it between HG and pmSTG (Figure 5). In other words, although a previous study showed that the auditory cortex had stronger connectivity with InsP than with InsA (Zhang et al., 2018), we argue that the functional connectivity between InsP and the primary auditory cortex is far weaker than the connectivity between primary and non-primary cortex. The current evidence suggests that InsP might be rather independent of the auditory cortical system and further support the multi-sensory integration view.

In summary, the data in the current study suggested that InsP could not be defined as a part of the auditory sensory cortex. Although the basic response properties of auditory evoked high-γ in InsP are similar to primary or non-primary auditory cortices in the supra-temporal plane, the InsP contribute little to the production of auditory sensation and may function as an early monitoring system for integrating auditory and somatosensory.

## METHODS

### Participants

Fifty-five patients suffering from drug-refractory epilepsy (25 females, aged from 9 to 46 years old, details in Table S1), who were recruited in the Sanbo Brain Hospital of Capital Medical University, participated in this study. The participants were undergoing long-term invasive sEEG monitoring to identify their seizure foci. During the monitoring period, all the participants were weaned from their antiepileptic medications. All participants were right-handed with a self-reported normal hearing and provided written informed consent for their participation. Thirty-two patients were implanted with sEEG in bilateral hemispheres, while 13 patients were implanted in the left hemisphere and 10 in the right hemisphere (Table S1). The experimental procedures were approved by the Ethics Committee of the Sanbo Brain Hospital of Capital Medical University.

### sEEG data acquisition and experimental procedures

Stereo-EEG signals were recorded using the Nicolet video-EEG monitoring system (Natus Neuro, USA), digitized at the rate of 512 Hz and collected with a 0.05-200 Hz online bandpass filter. The sEEG electrodes were manufactured by Huake Hengsheng Medical Technology Co Ltd, Beijing, China. The diameter of a depth electrode was 0.8 mm. The length of each contact was 2 mm, which were spaced 1.5 mm apart from each other. The reference electrode was placed on the forehead. On the experimental day, the impedance of all the recording contacts was kept below 50 kΩ.

Gaussian wideband-noise stimuli were synthesized in MATLAB environment (MathWorks, Natick, MA, USA) at the sampling rate of 48 kHz with 16-bit amplitude quantization and low-pass filtered at 10 kHz. The duration of the noise-burst stimulus was 50 ms including the 5-ms linear ramp and damp. The acoustic stimulus was transferred using Creative Sound Blaster (Creative SB X-Fi Surround 5.1 Pro, Creative Technology Ltd, Singapore) and presented to participants with insert earphones (ER-3, Etymotic Research, Elk Grove Village, IL) at the sound pressure level of 65 dB SPL. Calibration of the sound level was carried out with the Larson Davis Audiometer Calibration and Electroacoustic Testing System (AUDit and System 824, Larson Davis, Depew, NY).

Each participant was reclining on a bed-ward in the hospital during the experiment. The experimental procedure was suspended for at least 3 hours after a seizure to avoid the seizure-induced cortical suppression effect. Participants were asked to stay awake and passively listened to the noise bursts without any behavioral task. Nine patients were only presented with binaural sounds while 46 patients were presented with sounds from ipsilaterally, contralaterally and binaurally ears, respectively (Table S1). Each condition was repeated 320 or 160 trials. The inter-stimulus interval (ISI) between the two stimulus presentations was random between 900 to 1000 ms.

### Brain reconstruction and electrode contact localization

Three-dimensional brain surfaces were reconstructed by pre-implantation MR images (T1 or contrast-enhanced), registered using USC Brain atlas (Joshi et al., 2017) in BrainSuite (http://brainsuite.org/). Using BioImage (http://bioimagesuite.yale.edu) software, the original coordinates of electrode contacts were extracted from the images with the fusion between the pre-implantation MR and the post-implantation CT scans. The localization of sEEG electrodes was then presented using the Brainstorm toolbox (Tadel et al, 2011; http://neuroimage.usc.edu/brainstorm/) in the MATLAB environment. The localization of each contact was identified using an individual atlas of each patient. The contacts localized in the anterior insula (InsA), posterior insular (InsP), Heshl’s gyrus (HG), anterior superior temporal gyrus (aSTG), middle superior temporal gyrus (mSTG), and posterior superior temporal gyrus (pSTG), were selected. All the subject-wised coordinates were transferred into MNI coordinates for group plottings.

### sEEG data analyses

In each participant, epileptic foci had been identified before our experiment. The electrode contacts with over threshold impedances (> 50 kΩ), artifacts, and/or within epilepsy foci were excluded from data analyses.

All the electrophysiological signals were pre-processed using the EEGLAB toolbox (Delorme and Makeig, 2004) in the MATLAB environment. To extract the high-γ activity, the long-term signals were filtered by band-pass filters (60-140 Hz) and then translated into a power envelope via Hilbert transforms. The long-term high-γ energy was then segmented into epochs from -100 to 600 ms around the sound onset. The time window of the baseline was defined from -100 to 0 ms before the sound onset. The epochs which contained transient artifacts that marked in the broadband signal analysis were rejected. To decide whether an individual contact was significantly responsive to auditory stimuli, trial-level Student’s *t*-tests against baseline (with *Bonferroni* correction) were conducted for the time points after the sound onset. The contact with at least one significant point was identified as significantly auditory responsive and entered further analyses. The time of the earliest significant time point was extracted as the latency of the contact.

To compared the high-γ activations under contralateral and ipsilateral conditions, trial-level independent *t*-tests (with *Bonferroni* correction) within early (0 to 50 ms) and late (50 to 300 ms) time windows were calculated, respectively. The *t* value was extracted as the contralateral to ipsilateral (CI) index under each comparison.

### Unsupervised clustering

According to a previous study, the waveforms of auditory evoked high-γ can be divided into onset and sustained types (Hamilton et al., 2018). To verify this finding in the human supra-temporal and insular auditory responsive region, we also used a non-negative matrix factorization toolbox (Li and Ngom, 2013) to cluster the time series of the evoked high-γ activity. The time series X [time points × contacts] were estimated with the function (1):

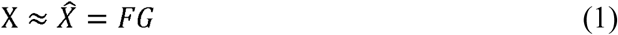

Where *G* matrix [clusters × contacts] represents the classification weighting of a contact on a certain cluster and *F* matrix [time points × clusters] represents the prototypical time series of a given cluster.

All the time series of significant high-γ contacts from all patients were considered as *X* [614 time points × 524 contacts]. To evaluate the explanatory power of the results, the percent variance (*R* square) was calculated with the function (2):

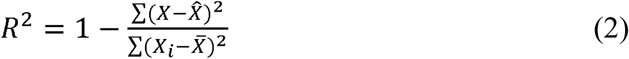

The *R* squares with different cluster numbers (from *k* = 2 to *k* =10) were calculated to evaluate the optimal cluster number (Figure S3). For the optimal cluster, the silhouette index (SI) was calculated to evaluate cluster separability using the *silhouette* function in Matlab under each condition. The classification types for all contacts were shuffled 1000 times to create the chance levels of SI for statistical tests.

### Granger causality analysis

Granger causality (GC) analyses (time domain) were calculated using the Brainstorm toolbox. The GC is considered from X to Y (i.e., X→Y) if including past values of X and Y (i.e., full model) provides more information about future values of Y compared to when only the past values of Y (i.e., restricted model) are considered (Seth, 2010). Here, X or Y is the time series representing auditory evoked high-γ responses with a time window from 0 to 300 ms in contact level. Note that contacts only from the same patient were paired and used for GC analyses to avoid individual differences from calculating. The higher GC value represents a stronger interaction from X to Y. To assess the statistical significance (*p*-value) of the GC value between two electrodes X→Y, we tested the null hypothesis (i.e., the full model did not fit the data better than the restricted model) using the *F*-statistic. Only the significant GC values were entered into further analyses.

### Cortico-cortical evoked potential (CCEP)

Four out of 55 patients participated in the CCEP experiment. Single-pulse electrical stimulations (pulse width = 0.2 ms, trial number = 20, amplitude = 2, 4, and 6 mA) were performed through the contacts which located in HG, InsP pSTG or mSTG via Nicolet® Cortical Stimulator (Natus Neuro, USA). The procedures were conducted at the bedside while the patients were awake and resting. An on-line 50 Hz notch-filter was applied to reduce the line-noise.

The results of CCEP mapping were further analyzed using Matlab codes. The evoked responses were divided into epochs from -100 ms to 600 ms according to the stimulation pulse onset and the bad trials were deleted according to both visual inspection and voltage threshold criteria. For that the stimulation artifact could be observed around the onset period, the baseline was set from -100 to -10 ms for drift corrections. Then the epochs under each condition were averaged to get the CCEP waveform. The strength of the CCEP amplitude was calculated using the root-mean-square value from 10 to 300 ms.

### Electrical brain stimulation (EBS)

Forty-two out of 55 patients participated in electrical brain stimulation (EBS) tests for cognitive function evaluation using Nicolet® Cortical Stimulator (Natus Neuro, USA). After the recordings of clinical seizures, electrical stimulations were performed through the electrodes in each contact pairs (frequency = 50 Hz, pulse width = 1 ms, duration = 5 seconds, amplitude from 1 to 6 mA). Patients were unaware of the timing of the stimulation and the anatomical location of the stimulated structure. During the period, patients were asked to performing a number counting task (speak loudly from one to one hundred). Patients were asked to report immediately if they had any feelings. The results of patients’ self-reported were reviewed and further classified by at least one experienced neuropsychologist.

### Micro-contact recordings using a hybrid electrode

In only 1 in 55 patients (P374), we recorded the extracellular potential in left HG and InsP using hybrid depth electrodes (2069-EPC-8C9-35T04, ALCIS Neuro, France) with 9 micro-contacts (20 µm diameter) interspersed between 4 of the 8 cylindrical macro-contacts (0.8 mm diameter; 2 mm length) (Figure 6A). The patient was a 46-year-old right-handed woman with 12 years history of seizures. Continuous intracranial EEG signals from all the micro contacts were recorded in an unshielded hospital room using a 256-channel NATUS Xltech digital video-EEG system (10–50 MΩ amplifier impedance; 16 kHz sampling rate). The procedures of auditory stimulus presenting the same as above (bilateral 50 ms noise burst, 160 trials).

To estimate the location of the electrode contacts, the post-implantation CT scan was aligned to the post-implantation MRI scan. The micro contacts were not visible on the CT scan but were situated between the macro contacts (Figure 6B).

The signals were off-line filtered by a band-pass filter (0.5-200 Hz) segmented into epochs from -200 to 1000 ms around the sound onset. The baseline correction was conducted by the time window from -200 to 0 ms before the sound onset. Signal trial *t*-tests (with *Bonferroni* correction) against the baseline were analyzed in the Brainstorm toolbox.

## Acknowledgments

We give our highest respect to all the patients who participated in this study. We thank all the neurologists, nurses and technicians at the Epilepsy Center of Sanbo Hospital who participated in the care of these patients and helped make this research possible. This work was supported by the National Natural Science Foundation of China (no. 31700994, 81790654, 81790650, 31871131).

