## Supplemental Figure 1-4, Table 1-2 for "Revealing the functions of supra-temporal and insular auditory responsive areas in humans": SuppM200330.pdf

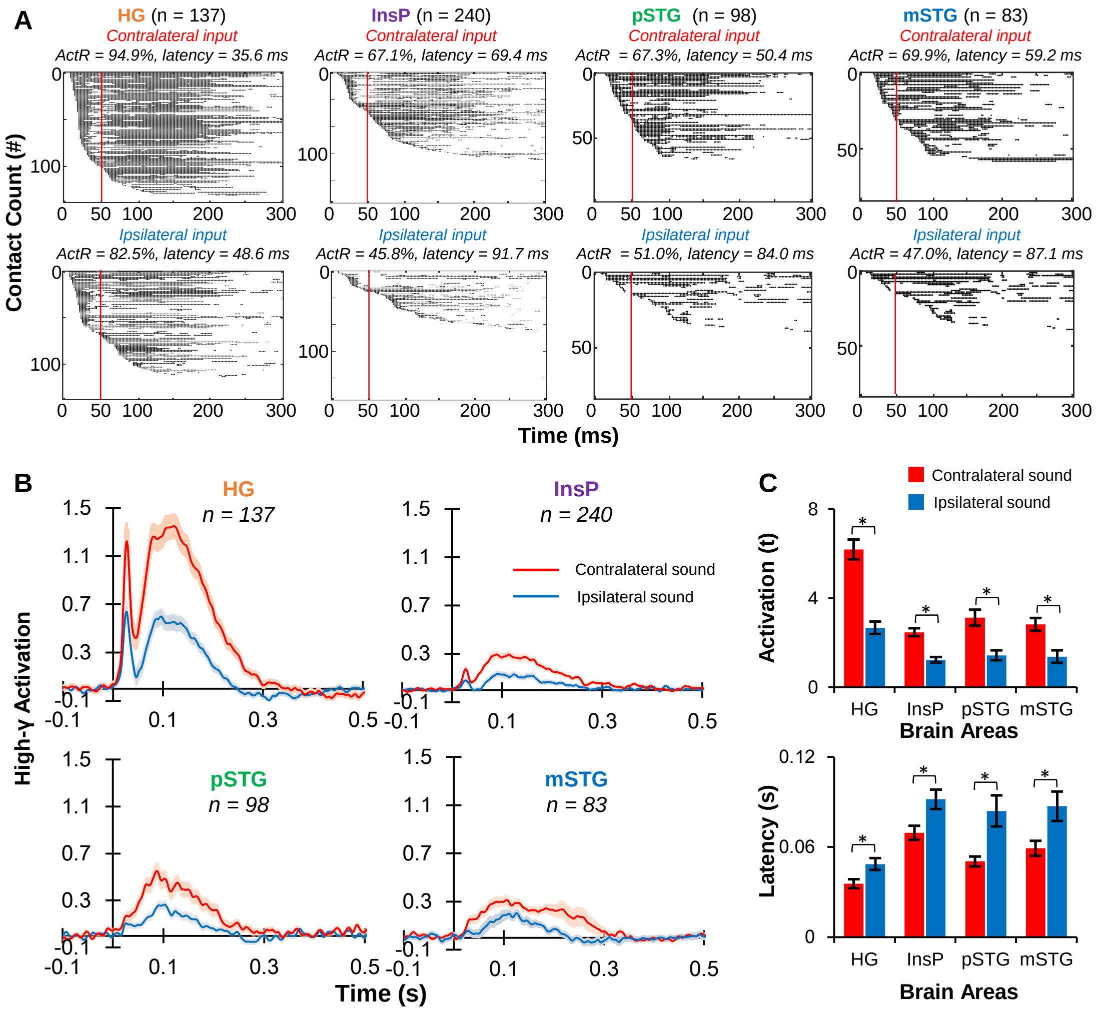


**Figure S1. The contralateral input dominance in the human supra-temporal area and InsP.** (A) The significance map of the auditory evoked high-γ within individual contact in HG, InsP, pSTG, and mSTG. Each line represents one contact and black dots represent the significant time points against the baseline (*t*-tests in single-trial level, *Bonferroni* corrected $p < 0.05$). (B) The averaged high-γ responses under contralateral and ipsilateral input conditions. Shaded areas indicate standard errors. (C) Comparison of the activations and latencies between contralateral and ipsilateral conditions within each ROI. *, $p < 0.05$.

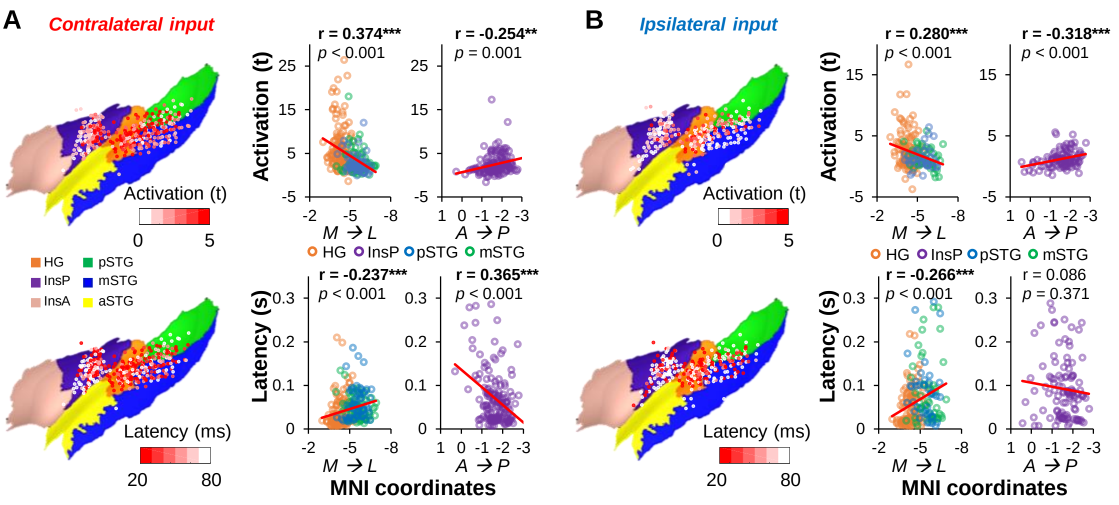


**Figure S2. The distributions of activation and latency under contralateral and ipsilateral conditions.** Contacts located on HG, InsP, pSTG, and mSTG are color-coded by activations and latencies. Scatter plots show how activations and latencies are correlated with their MNI coordinates (Pearson correlation tests). (A) Contralateral input condition. (B) Ipsilateral input condition. *A→P*, from anterior to posterior; *M→L*, from medial to lateral. **, $p < 0.01$; ***, $p < 0.001$.

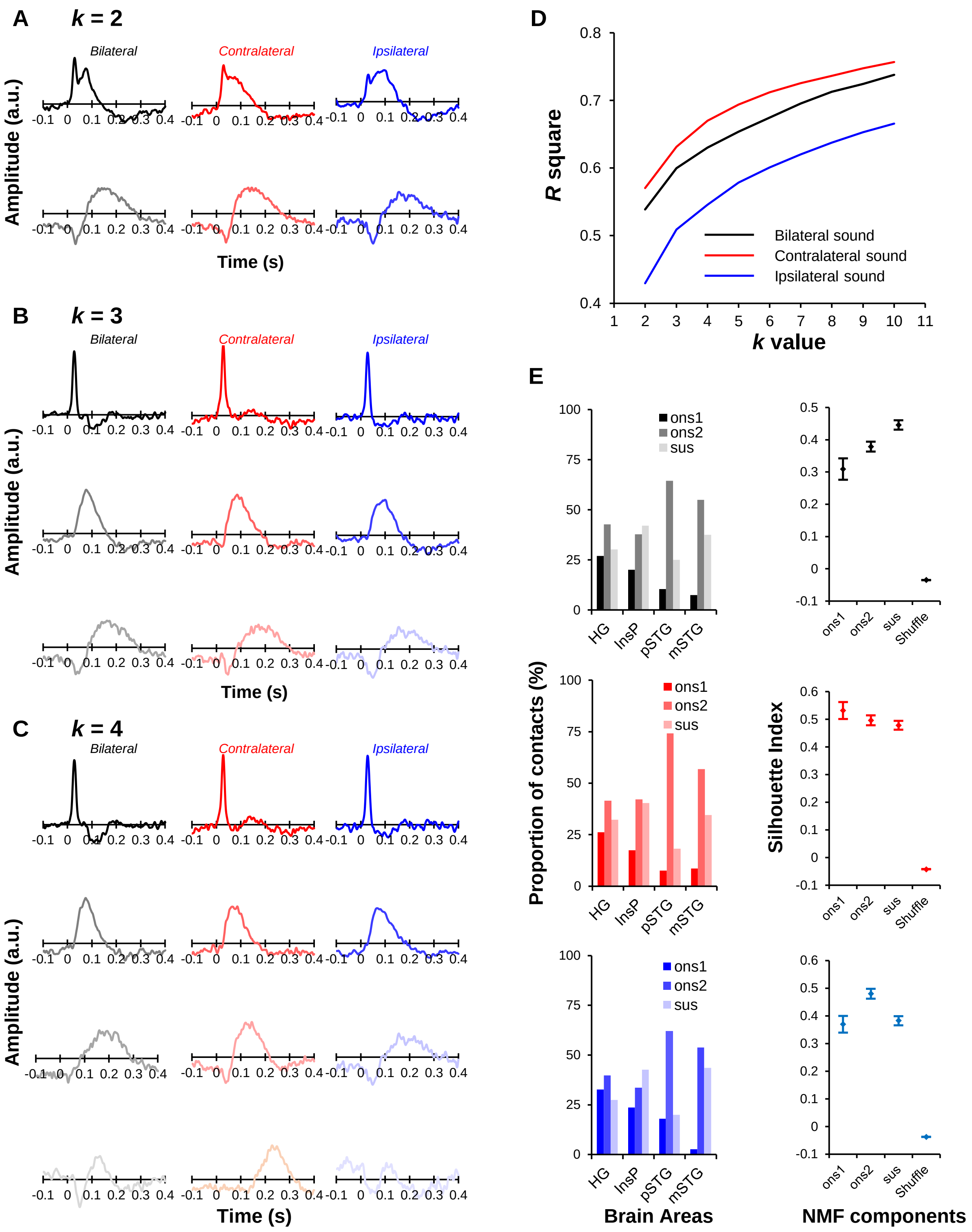


**Figure S3. Unsupervised clustering results for different numbers of clusters (k) under three stimulation conditions, related to Figure 2.** (A), (B), and (C) show the prototypical time series of clustering results for k =2, 3, and 4, respectively. In (A), (B) and (C), the clustering results under three stimulation conditions are similar. (D) Explanatory powers (R square) of cluster numbers from 2 to 10. The increasement goes slow when the clusters are beyond 3. (E) Proportion of the contact among 4 ROIs and the silhouette index (SI) under $k$ = 3 conditions. All SIs were significantly larger than the shuffled level (all $p < 0.001$, paired $t$-tests).

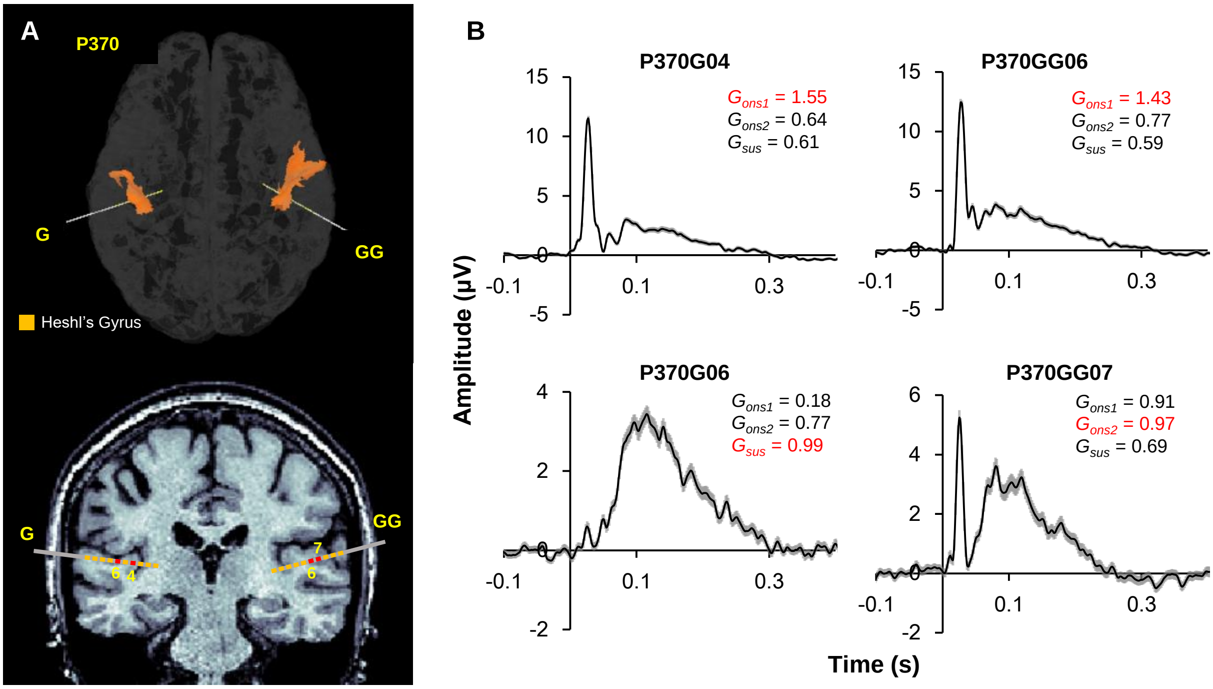


**Figure S4. Illustration of weight-depended classification for individual contact in P370.** (A) Location of a pair of 10-contact sEEG electrodes implanted on the left (electrode G) and right (electrode GG) HG, respectively. (B) Averaged sound-induce high-γ responses (bilateral input condition) from 4 HG contacts and their classification weights for three clusters (Gons1, Gons2, and Gsus). Red fonts show the corresponding classification for each contact. Shaded areas indicate standard errors.

**Supplementary Table 1. Patients' demographic and clinical data**

| Patient# | Age | Sex | Implanted Hemisphere | Contacts Number | Selected Hemisphere | Selected Contacts | Insula | HG | STG |
|---|---|---|---|---|---|---|---|---|---|
| 121 | 24 | F | L | 105 | L | 20 | 9 | 5 | 6 |
| 161 | 17 | F | L, R | 117 | L | 13 | 7 | 3 | 3 |
| 172 | 30 | F | L, R | 115 | R | 12 | 8 | 1 | 3 |
| 206 | 26 | M | L, R | 117 | L | 15 | 5 | 7 | 3 |
| 207 | 28 | M | L, R | 124 | R | 23 | 17 | 3 | 3 |
| 208 | 17 | M | L, R | 121 | L | 17 | 10 | 3 | 4 |
| 220 | 26 | M | L | 111 | L | 11 | 8 | 3 | 0 |
| 229 | 28 | M | R | 123 | R | 10 | 2 | 4 | 4 |
| 233 | 26 | M | L, R | 120 | R | 9 | 3 | 3 | 3 |
| 235 | 24 | F | L, R | 112 | L | 6 | 2 | 2 | 2 |
| 236 | 18 | M | L, R | 112 | R | 22 | 18 | 2 | 2 |
| 237 | 27 | F | L | 121 | L | 16 | 9 | 2 | 5 |
| 240 | 29 | F | L | 122 | L | 13 | 5 | 2 | 6 |
| 243 | 27 | M | L, R | 113 | R | 8 | 2 | 3 | 3 |
| 245 | 31 | F | L, R | 116 | R | 10 | 10 | 0 | 0 |
| 260 | 36 | F | L, R | 120 | L | 10 | 5 | 3 | 2 |
| 263 | 28 | M | L | 117 | L | 6 | 1 | 3 | 2 |
| 272 | 31 | M | L, R | 121 | R | 8 | 2 | 3 | 3 |
| 276 | 20 | M | L, R | 116 | L | 21 | 11 | 5 | 5 |
| 277 | 26 | F | L, R | 121 | R | 10 | 4 | 0 | 6 |
| 279 | 33 | F | L, R | 126 | L, R | 35 | 24 | 2 | 9 |
| 280 | 23 | M | L, R | 108 | L, R | 30 | 13 | 6 | 11 |
| 281 | 31 | F | L, R | 122 | L | 15 | 11 | 4 | 0 |
| 282 | 27 | F | L, R | 124 | L | 13 | 8 | 5 | 0 |
| 290 | 26 | M | L, R | 122 | L, R | 31 | 27 | 2 | 2 |
| 358 | 38 | M | L, R | 120 | R | 10 | 2 | 2 | 6 |
| 359 | 36 | F | R | 118 | R | 25 | 17 | 2 | 6 |
| 361 | 27 | F | R | 118 | R | 16 | 13 | 3 | 0 |
| 364 | 27 | M | L | 126 | L | 29 | 17 | 5 | 7 |
| 365 | 27 | M | L, R | 116 | L | 21 | 6 | 4 | 11 |
| 370 | 25 | F | L, R | 128 | L, R | 41 | 28 | 8 | 5 |
| 371 | 24 | M | L, R | 116 | L | 9 | 8 | 1 | 0 |
| 373 | 15 | M | R | 114 | R | 18 | 10 | 5 | 3 |
| 374 | 46 | F | L, R | 122 | L | 26 | 15 | 2 | 9 |
| 377 | 45 | M | L, R | 124 | L | 22 | 7 | 6 | 9 |
| 380 | 17 | F | R | 116 | R | 11 | 2 | 2 | 7 |
| 381 | 27 | F | L | 119 | L | 33 | 21 | 4 | 8 |
| 383 | 17 | M | L | 105 | L | 13 | 2 | 2 | 9 |
| 384 | 14 | F | R | 108 | R | 18 | 5 | 2 | 11 |
| 386 | 29 | M | L, R | 128 | L, R | 25 | 8 | 2 | 15 |

| | | | | | | | | | |
|---|---|---|---|---|---|---|---|---|---|
| 387 | 9 | F | R | 110 | R | 24 | 14 | 2 | 8 |
| 389 | 19 | M | R | 122 | R | 13 | 7 | 2 | 4 |
| 390 | 26 | M | L | 100 | L | 5 | 5 | 0 | 0 |
| 392 | 13 | M | L | 124 | L | 25 | 14 | 3 | 8 |
| 409 | 21 | M | R | 118 | R | 14 | 3 | 3 | 8 |
| 412 | 29 | M | L | 124 | L | 12 | 4 | 3 | 5 |
| 413 | 43 | F | L, R | 118 | L | 23 | 12 | 3 | 8 |
| 414 | 31 | M | L, R | 124 | L | 16 | 4 | 4 | 8 |
| 417 | 35 | F | R | 124 | R | 15 | 9 | 3 | 3 |
| 419 | 34 | N | L, R | 128 | R | 12 | 2 | 6 | 4 |
| 420 | 46 | F | L, R | 128 | L | 19 | 3 | 6 | 10 |
| 421 | 13 | F | L | 126 | L | 21 | 14 | 2 | 5 |
| 422 | 22 | M | L | 126 | L | 26 | 12 | 3 | 11 |
| 430 | 31 | F | L, R | 112 | L, R | 25 | 7 | 1 | 17 |
| 435 | 24 | M | L, R | 126 | L | 2 | 1 | 1 | 0 |
| Sum | | | | | | 953 | 493 | 168 | 292 |

F, female; M, male; L, left; R, right; HG, Heshl’s Gyrus; STG, superior temporal gyrus.

**Supplementary Table 2. Proportion of different EBS evoked sensation in four ROIs.**

| | **Auditory Suppression** | **Auditory Hallucination** | **Noxious Sensation (burring, stinging, electrical shock)** | **Somatosensory Sensation (warmth, cooling)** | **Epigastric Sensation** | Vestibular Reponses | Speech Arrest | **Illusion** |
|---|---|---|---|---|---|---|---|---|
| ***HG*** | 9 | 80 | 6 | 6 | 1 | 4 | 3 | 0 |
| (102/133) | (8.82%) | (78.43%) | (5.88%) | (5.88%) | (0.98%) | (3.92%) | (2.94%) | (0.00%) |
| ***InsP*** | 0 | 15 | 32 | 109 | 6 | 3 | 0 | 3 |
| (158/221) | (0.00%) | (9.49%) | (20.25%) | (68.99%) | (3.80%) | (1.90%) | (0.00%) | (1.90) |
| ***pSTG*** | 1 | 18 | 0 | 3 | 0 | 5 | 0 | 0 |
| (27/63) | (3.70) | (66.67) | (0.00%) | (11.11%) | (0.00%) | (18.52%) | (0.00%) | (0.00%) |
| ***mSTG*** | 3 | 21 | 2 | 5 | 0 | 3 | 0 | 1 |
| (28/71) | (10.71%) | (75%) | (7.14%) | (17.86%) | (0.00%) | (10.71%) | (0.00%) | (3.57%) |

** (contacts with positive sensation/ all contacts); EBS, electrical brain stimulation; ROI, region of interest.*
